# Comparison of Analytical Techniques for Thermal Stability Analysis of β-Cyclodextrin for an Ebola Virus Infection Treatment

**DOI:** 10.1101/448928

**Authors:** Mayte Murillo, Robert A. Lodder

## Abstract

Each New Drug Application filed with the Food and Drug Administration (FDA) must include the analytical procedures to ensure the identity, strength, quality, purity, and potency of a drug substance and drug product. The BSN389 drug product (being developed to treat Ebola virus infections) includes beta cyclodextrin. Evidence must be provided that the analytical procedures used in testing BSN389 meet proper standards of accuracy, sensitivity, specificity, and reproducibility and are suitable for their intended purpose. The Bootstrap Error-adjusted Single-Sample Technique (BEST) software was used to compare the quantitative and qualitative power of IR and ^1^H NMR to differentiate new and partially decomposed samples of beta cyclodextrin, and the best assay will be incorporated into the thermal stability protocol for ВCD.

## 1. INTRODUCTION

Beta cyclodextrin, known simply as β-cyclodextrin or βCD or ВCD, is a non-reducing cyclic oligosaccharide consisting of seven α-1,4- linked D-(+)-glucopyranosyl units (Figure 1). The seven membered ring is produced by enzymatic conversion of starch. This drug has applications not only in pharmaceuticals but also in the food and environmental industry. Toxins can be removed when the ring ensnares specific molecules that are targeted for removal. ВCD is also a food additive that acts as a stabilizer for flavors, colors, and some vitamins.^2^ ВCD’s estimated intake is about 1-1.4 g/day and it is approved by the FDA. Researchers are nowtaking known information about the cyclodextrin molecule and using it as a carrier for chemotherapeutic cytotoxic anticancer drugs.^3^

**Figure 1.**
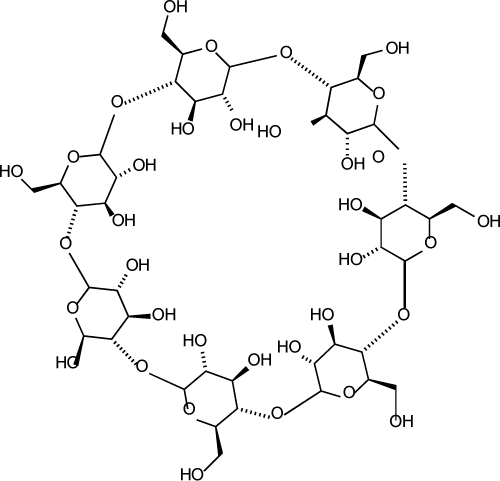
Structure of β-cyclodextrin.

The analytical techniques ^1^H NMR and infrared (IR) spectro––scopy are used to measure the difference between decomposed and stable versions of ВCD in this research. The Bootstrap Error-adjusted Single-sample Technique (BEST) software (see Appendix) will then identify the best analytical method to use for the thermal stability regulatory procedure for the drug.

## 2. METHODS

### Preparation of Samples

Approximately two grams of beta cyclodextrin was slightly decomposed thermally by putting the sample on a Pyrex dish and placing it in a conventional oven, heating it slowly to about 232 °C at a linear rate of about 28 °C/5 min until the white powder sample was a slightly yellow color.

#### Measurements

Six separate samples of the pure and decomposed ВCDs were prepared with deuterated water. H NMR spectra were recorded on a 500 MHz JOEL spectrometer, and processed with 16 scans ranging from −2 to 16 ppm. These samples were then also analyzed using a Thermo Scientific Nicolet iS10 infrared spectrometer over a wavenumber range of 4000-500 cm^-1^.

#### Analysis

The ^1^H NMR data were entered in TopSpin and converted into CVS files. These data along with the IR values were read into MATLAB. Each sample set of data was linked together in a variable with dimensions equal to the number of wavenumbers or chemical shifts. Each set was plotted as described in Appendix 1. The BEST program was used to determine the distance in multidimensional standard deviations (SDs) between the set of samples of pure and decomposed ВCD. These values were then compared to the control distances, which were found by finding the distances between the center of the pure ВCD validation spectra each pure ВCD validation spectrum.

### 3 RESULTS AND DISCUSSION

#### Characterization of ВCD

As shown in Figure 2 and 3, the structure of ВCD was characterized by^1^H-NMR and IR spectroscopy. Figure 2 shows the^1^H-NMR spectrum of decomposed and pure ВCD. Results showed only slight left shift of the decomposed sample. Figure 3 shows a comparison of the IR spectra between the pure and decomposed drug. The decomposed sample showed a slight blue shift in the O-H stretch vibrations.

**Figure 2.**
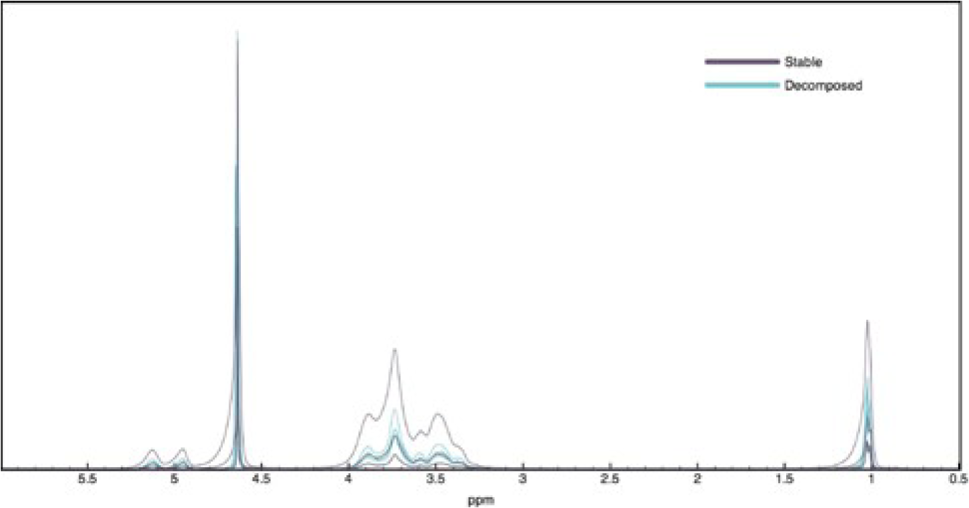
^1^H NMR forВCD

**Figure 3.**
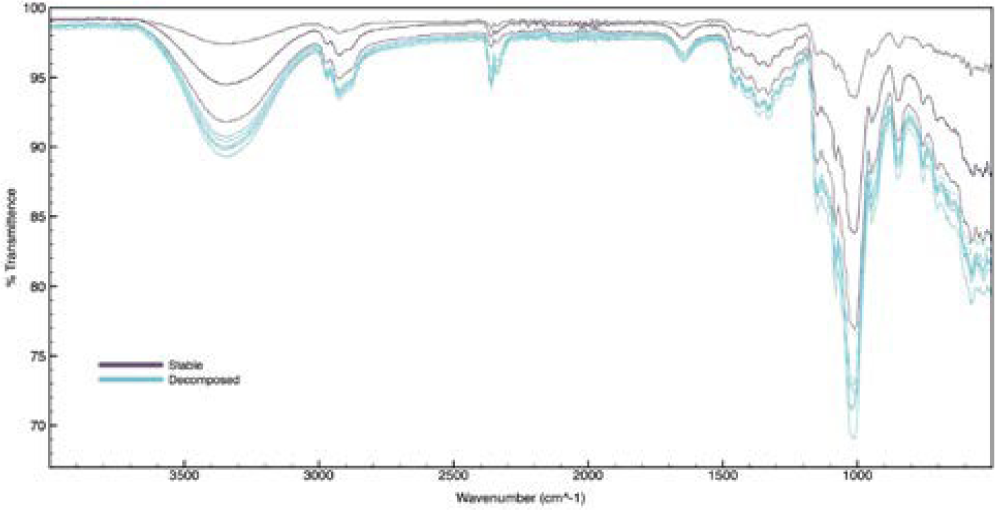
IR spectra of ВCD

#### Analysis of Techniques

Table 1 shows the standard deviations output by the BEST software. The average values for the^1^H NMR were higher than the average values for the IR, 6.3100 and 4.9900 respectively. This showed the differences between the stable and decomposed sample that are not obvious when shown graphically (Figures 2 and 3). The controls were expected to be less than three. There were two outliers in the NMR spectra. The outliers will be checked by LCMS in a follow-up study.

**Table 1.** BEST results comparing pure and decomposed samples.

| Sample | NMR |  | IR |  |
| --- | --- | --- | --- | --- |
|  | Pure/Decomposed | Pure/Pure | Pure/Decomposed | Pure/Pure |
| 1 | 3.7103 | 1.8613 | 4.3276 | 1.2500 |
| 2 | 3.6602 | 1.8611 | 4.8218 | 1.5973 |
| 3 | 1.6542 | 1.6542 | 4.7514 | 1.7374 |
| 4 | 11.6069 | 0.9857 | 5.0882 | 1.8101 |
| 5 | 10.6345 | 2.1662 | 5.8849 | 1.3649 |
| 6 | 6.5937 | 2.3055 | 5.0665 | 1.4278 |

### 4. CONCLUSION

Beta cyclodextrin samples were tested by IR and NMR. Tests showed small differences between the pure and slightly decomposed samples. The BEST software suggests that proton NMR is superior for thermal stability of ВCD, but the variability is large and the outliers need to be investigated.

It is worth noting that the Mahalanobis distance between the IR and NMR samples could not be calculated because the number of rows of the data matrix must exceed the number of columns. The BEST software was able to overcome this obstacle and provide a useful result.

## 5. ACKNOWLEDGEMENTS

The project described was supported by the National Center for Research Resources and the National Center for Advancing Translational Sciences, National Institutes of Health, through Grant UL1TR001998, and by the National Science Foundation through NSF ACI-1053575 allocation number BIO170011. The content is solely the responsibility of the authors and does not necessarily represent the official views of the NIH.

## APENDIX I

~~~
****REPLICA PROGRAM****
function [BTRAIN,CNTER]=replica(TNSPEC,B)
% TNSPEC=training spectra, B=number of replicates desired.
% REPLICA outputs BTRAIN replicates, and the center of the replicates in CNTER
%“Copyright 2003 Robert A. Lodder & Bin Dai”
     [N,D]=size(TNSPEC);
     BTRAIN=zeros(B,D);
     CNTER=zeros(D,1);
     BSAMP=zeros(N,D);
     PICKS=rand(B*N,1);
     index=find(PICKS==1);
     PICKS(index)=0.9999;
     PICKS=reshape(PICKS,B,N);
     PICKS=fix(N*PICKS+1);
     for I=1:B
          BSAMP=TNSPEC(PICKS(I,:),:);
          BTRAIN(I,:)=sum(BSAMP)/N;
     end
     BTRAIN;
     CNTER=sum(BTRAIN)/B;
****QB PROGRAM****
% Copyright 2003 Robert A. Lodder
function [sds,sdskew] = qb(tnspec,btrain,newspec,cnter,radfrac,sensitiv)
% function definition of qb with parameters
% tnspec= which is training spectra
% btrain =bootstrap replicates calculated using routine replica
% newspec= sample spectrum
% cnter= center of calibration set calculated using routine replica
% radfrac= fraction of points in hypercylinder
% sensitiv= sensitivity
b = size(btrain,1);     % finds the number of rows in btrain
qdist = zeros(1,b);     % creates a null row matrix
s02 = sqrt(sum((newspec-cnter).^2));     % computing the squareroot of sum of the suares
s0r = sqrt(sum((btrain-repmat(cnter,b,1)).^2,2)); % repmat creates a bX1 tilings of cnter
s2r = sqrt(sum((btrain-
repmat(newspec,b,1)).^2,2)); sub =
(s02+s0r+s2r)/2;
area = sqrt(sub.*(sub-s02).*(sub-s0r).*(sub-s2r));
radial = (2*area)/s02;
project = sqrt(s0r.^2-radial.^2);          % finds the indices where s02*s02 + s0r*s0r
locs = find((s02.^2+s0r.^2) < s2r.^2);
< s2r*s2r(getting the locus)
project(locs) = project(locs)*-1;
qdist = project;
qrr = sort(radial);     %sorts the elements in ascending order
radii = qrr(radfrac*b);
qdist(find(radial > radii)) = 0; % setting the elments of qdist to zero where
radi-
al > radii
qdist = sort(qdist(find(qdist))); % sorts all non zero elememts in qdist
lindex = floor(0.16*length(qdist)); % lower limit found by rouding to nearest
inte-ger
uindex = floor(0.84*length(qdist)); % upper limit found by rounding to nearest
in-teger
if(length(qdist) < 50)
’** Need more replicates in hypercylinder **’
end
sd = std(qdist)*sqrt(size(tnspec,1)); % std returns the standard deviation
sds = sqrt(sum((cnter-newspec).^2))/sd;     % calculation of the standard
deviation distances
% BIAS ADJUSTMENT
alpha = normcdf(−1,0,1); % computes the cumulative distribution function with mean 0 and standard deviation 1
za = norminv(alpha,0,1); % inverse of the cumulative ditribution function with mean o and standard deviation 1
tcenter = median(tnspec); % finding the
median cs02 = s02;
cs0r = sqrt(sum((tcenter-cnter).^2));
cs2r = sqrt(sum((tcenter-newspec).^2));
csub = (cs02+cs0r+cs2r)/2;
carea = sqrt(csub*(csub-cs02)*(csub-cs0r)*(csub-
cs2r)); cradial = (2*carea)/cs02;
cproject = sqrt(cs0r^2-cradial^2);
if((s02^2+cs0r^2) > cs2r^2)
          cproject =-cproject;
end
n = length(qdist); % finds the length of the vector
if(floor(n/2) == n/2)
          md = (qdist(n/2)+qdist(n/2+1))/2;
else
          md = qdist(floor(n/2+0.5));
end
cproject = cproject*sensitiv + md;
fdist = qdist-cproject;
index = 1:length(fdist);
if(cproject > max(qdist))
          zelement = length(qdist)-1;
elseif(cproject < min(qdist))
          zelement = 1;
else
          rootloc =
          find(abs(fdist)==min(abs(fdist)));
zelement = rootloc(1);
end
z0 = norminv(zelement/length(qdist),0,1);
if(abs(2*z0) > abs(za))
          error(’ –– Decrease skew sensitivity. ––’);
end
sensitiv = abs(sensitiv);
lowind =
floor(normcdf(2*z0+za,0,1)*length(qdist)); upind
= floor(normcdf(2*z0-za,0,1)*length(qdist));
if(lowind < 2)
         ‘** Warning ** Too few replicates’
end
if(upind > length(qdist)-2)
         ‘** Warning ** Too few replicates’
end
if(lowind < 1)
          lowind = 1;
end
if(upind > length(qdist))
          upind = length(qdist);
end
lowlim = qdist(lowind);
uplim = qdist(upind);
euc = sqrt(sum((cnter-newspec).^2));
fac = abs(norminv(alpha));
erd = sqrt(size(tnspec,1));
if(abs(2*z0)>abs(fac))
         ‘** Warning ** SKEW CORRECTION exceeds replicates’
end
sdskew = euc/((uplim/fac)*erd);
~~~

~~~
**NMR COMPARISON**
>> NMR_stable=zeros(2,2);
NMR_decomposed=zeros(2,2);
>> ppm=zeros(2,2);
>> plot(ppm,NMR_stable);
hold on
plot(ppm,NMR_decomposed);
hold off
[BTRAIN,CNTER]=replica(NMR_stable,1000);
[sds,sdskew] = qb(NMR_stable,BTRAIN,NMR_decomposed(1,:),CNTER,0.25,0);
>> sds
sds =
3.7103
>> [sds,sdskew] = qb(NMR_stable,BTRAIN,NMR_decomposed(2,:),CNTER,0.25,0);
>> sds
sds =
3.6602
>> [sds,sdskew] = qb(NMR_stable,BTRAIN,NMR_decomposed(3,:),CNTER,0.25,0);
>> sds
sds =
1.6542
>> [sds,sdskew] = qb(NMR_stable,BTRAIN,NMR_decomposed(4,:),CNTER,0.25,0);
>> sds
sds =
11.6069
>> [sds,sdskew] = qb(NMR_stable,BTRAIN,NMR_decomposed(5,:),CNTER,0.25,0);
>> sds
sds =
10.6345
>> [sds,sdskew] = qb(NMR_stable,BTRAIN,NMR_decomposed(6,:),CNTER,0.25,0);
>> sds
sds =
6.5937
>> [sds,sdskew] = qb(NMR_stable,BTRAIN,NMR_stable(1,:),CNTER,0.25,0);
>> sds
sds =
1.8613
>> [sds,sdskew] = qb(NMR_stable,BTRAIN,NMR_stable(2,:),CNTER,0.25,0);
>> sds
sds =
1.8611 >>
[sds,sdskew] = qb(NMR_stable,BTRAIN,NMR_stable(3,:),CNTER,0.25,0);
>> sds
sds =
1.6542
>> [sds,sdskew] = qb(NMR_stable,BTRAIN,NMR_stable(4,:),CNTER,0.25,0);
>> sds
sds =
0.9857
>> [sds,sdskew] = qb(NMR_stable,BTRAIN,NMR_stable(5,:),CNTER,0.25,0);
>> sds
sds =
2.1662
>> [sds,sdskew] = qb(NMR_stable,BTRAIN,NMR_stable(6,:),CNTER,0.25,0);
>> sds
sds =
2.3055
~~~

## IR COMPARISON

~~~
>> IR_stable=zeros(2,2);
IR_decomposed=zeros(2,2);
wavenumber=zeros(2,2);
>> plot(wavenumber,IR_stable);
hold on
plot(wavenumber,IR_decomposed);
hold off
[BTRAIN,CNTER]=replica(IR_stable,1000);
>> [sds,sdskew] = qb(IR_stable,BTRAIN,IR_decomposed(1,:),CNTER,0.25,0);
>> sds
sds =
4.3276
>> [sds,sdskew] = qb(IR_stable,BTRAIN,IR_decomposed(2,:),CNTER,0.25,0);
>> sds
sds =
4.8218
>> [sds,sdskew] = qb(IR_stable,BTRAIN,IR_decomposed(3,:),CNTER,0.25,0);
>> sds
sds =
4.7514
>> [sds,sdskew] = qb(IR_stable,BTRAIN,IR_decomposed(4,:),CNTER,0.25,0);
>> sds
sds =
5.0882
>> [sds,sdskew] = qb(IR_stable,BTRAIN,IR_decomposed(5,:),CNTER,0.25,0);
>> sds
sds =
5.8849
>> [sds,sdskew] = qb(IR_stable,BTRAIN,IR_decomposed(6,:),CNTER,0.25,0);
>> sds
sds =
5.0665
>> [sds,sdskew] = qb(IR_stable,BTRAIN,IR_stable(6,:),CNTER,0.25,0);
>> sds
sds =
1.4278
>> [sds,sdskew] = qb(IR_stable,BTRAIN,IR_stable(5,:),CNTER,0.25,0);
>> sds
sds =
1.3649
>> [sds,sdskew] = qb(IR_stable,BTRAIN,IR_stable(4,:),CNTER,0.25,0);
>> sds
sds =
1.8101
>>[sds,sdskew] = qb(IR_stable,BTRAIN,IR_stable(3,:),CNTER,0.25,0); >> sds
sds =
1.7374
>> [sds,sdskew] = qb(IR_stable,BTRAIN,IR_stable(2,:),CNTER,0.25,0);
>> sds
sds =
1.5973
>> [sds,sdskew] = qb(IR_stable,BTRAIN,IR_stable(1,:),CNTER,0.25,0);
>> mahal(IR_decomposed(1,:),IR_stable)
Error using mahal (line 38)
The number of rows of X must exceed the number of columns.
~~~

## REFERENCES

(1) Solutions, N. C. Parchem - fine & specialty chemicals is a Leading Supplier of Enzymes such as Beta Cyclodextrin https://www.parchem.com/news-articles/Parchem-fine-specialty-chemicals-is-a-Leading-Supplier-of-Enzymes-such-as-Beta-CyclodextrinN000192.aspx" (accessed Dec 12, 2017).

(2) (2) EFSA PanelonFoodAdditivesandNutrient Sourcesadded toFood (ANS); Mortensen, A.; Aguilar, F.; Crebelli, R.; Di Domenico, A.; Dusemund, B.; Frutos, M. J.; Galtier, P.; Gott, D.; Gundert-Remy, U.; Leblanc, J.; Lindtner, O.; Moldeus, P.; Mosesso, P.; Parent-Massin, D.; Oskarsson, A.; Stankovic, I.; Waalkens-Berendsen, I.; Woutersen, R. A.; Wright, M.; Younes, M.; Boon, P.; Chrysafidis, D.; Gürtler, R.; Tobback, P.;Arcella, D.;Rincon, A.M.;Lambré, C. EFSA Journal2016, 14(12).

